# A Viral-Encoded Homologue of IPS-1 Modulates Innate Immune Signaling During KSHV Lytic Replication

**DOI:** 10.1101/2024.07.24.604690

**Authors:** Daniel Miranda, Buyuan He, Julio C. Sanchez, Ashkon Sennatti, Johnny R. Bontemps, James T. Tran, Wilson S. Tang, David Jesse Sanchez

## Abstract

Modulation of innate immunity is critical for virus persistence in a host. In particular, viral-encoded disruption of type I interferon, a major antiviral cytokine induced to fight viral infection, is a key component in the repertoire of viral pathogenicity genes. We have identified a previously undescribed open reading frame within the Kaposi’s sarcoma-associated herpesvirus (KSHV) genome that encodes a homologue of the human IPS-1 (also referred to as MAVS) protein that we have termed viral-IPS-1 (v-IPS-1). This protein is expressed during the lytic replication program of KSHV, and expression of v-IPS-1 blocks induction of type I interferon upstream of the TRAF3 signaling node including signaling initiated via both the RLR and TLR3/4 signaling axes. This disruption of signaling coincides with destabilization of the cellular innate signaling adaptors IPS-1 and TRIF along with a concatenate stabilization of the TRAF3 protein. Additionally, expression of v-IPS-1 leads to decreased antiviral responses indicating a blot to type I interferon induction during viral infection. Taken together, v-IPS-1 is the first described viral homologue of IPS-1 and this viral protein leads to reprogramming of innate immunity through modulation of type I interferon signaling during KSHV lytic replication.

## INTRODUCTION

Herpesviruses represent a significant cause of infectious morbidity in our society. Spread by seemingly innocuous sharing of fomites or saliva exchange, herpesviruses effectively establish persistent infection [1]. However, the need for control of herpesviral replication and pathogenesis is underscored by their manifestation of severe pathologies in immunocompromised individuals such as patients living with acquired immunodeficiency syndrome (AIDS). A significant percentage of the U.S. population is seropositive for KSHV (also called Human Herpesvirus-8 (HHV-8)), a virus identified in the mid-1990s as the etiologic agent of Kaposi’s Sarcoma, a disease with a characteristic pathology including severe, inflammatory linked skin lesions [2].

Pulmonary spread of these cancer-like lesions is ultimately fatal if the virus is left unchecked. In addition, KSHV has been linked to several lymphoproliferative diseases such as Body Cavity Based Lymphoma and Multicentric Castleman’s disease. In immunocompromised individuals these diseases represent a source of significant morbidity and mortality [3]. Infection by Kaposi’s Sarcoma-associated Herpesvirus (KSHV) and the resulting pathologic consequences continue to be a defining opportunistic infection in individuals living with AIDS.

Herpesviral diseases often manifest upon the severe disruption of the adaptive immune response in persons with AIDS. However, this begs to ask why the innate immune system cannot contain such an infection. Consequently, KSHV infection, as with any other infection, can only replicate effectively in a host if the virus encodes measures to counteract the host innate immune response. Invariably, human viral pathogens have at least one mechanism to deal with the antiviral type I interferon (IFN) system of innate immunity [4; 5]. KSHV, as with most herpesviruses, encodes several proteins that seem to modulate the IFN system [6; 7; 8; 9; 10]. The IFN system should allow for rapid antiviral responses against most infection – and thus presents itself as a critical system to understand and harness to fight against chronic infections like KSHV that seem to be well controlled normally by the immune system.

Modulation of innate immunity by chronic viruses including KSHV and HIV has been the focus of many groups over the past decade. Previously, our group described direct targeting of cellular IPS-1 by HIV infection to dampen IFN-based innate immune responses by HIV infection [11]. IPS-1 (IFN-beta promoter stimulator protein 1), also called MAVS (mitochondrial antiviral signaling protein), Cardif (CARD adaptor inducing IFN-beta), or VISA (virus-induced signaling adaptor), is a key component of inducing IFN after detection of cytoplasmic viral RNA through the RIG-I-like Receptor (RLR) pathway. Targeting of IPS-1 is critical for pathogenic viruses with several other viruses targeting IPS-1 ([12; 13; 14; 15; 16]). Here we identify a previously undescribed KSHV Open Reading Frame (ORF) we term viral-IPS1 (v-IPS-1). We show that this novel viral ORF is able to block IFN induction from multiple innate immune pathways and also impacts the stability of important proteins in these pathways.

## RESULTS

### KSHV Encodes a Homologue of IPS-1

To search for viral open reading frames with homology to IPS-1, we used BLAST to search through all of the DNA viral genome in the NCBI database (Figure 1A). Using BLAST, we translated all regions of the genomic DNA in different directions to find both characterized and uncharacterized ORFs. We found a predicted, but as of yet undescribed ORF in the genome of KSHV. Due to the similarity to cellular IPS-1, we named this new ORF viral IPS-1 (v-IPS-1) (Figure 1B). Viral IPS-1 contains homology in the IPS-1 proline rich domain as well as a region containing a TIM (TRAF3 interaction Motif). The central, proline rich domain of IPS-1 has been shown by previous work to contain a TRAF3-interaction motif (TIM) that functions in the direct interaction between IPS-1 and TRAF3 [17]. That interaction is required for proper IPS-1-mediated signaling. Because of the lack of CARD domains but retention of the PRO and TIM in v-IPS-1, our hypothesis is that interaction of v-IPS-1 and TRAF3 would disrupt the normal IPS-1/TRAF3 signaling thus crippling IFN induction converging from multiple innate immune receptors (e.g. RLRs or TLR3/4). Of note is that no homologous domain in the amino-terminal region of v-IPS-1 that would correspond to the IPS-1 CARD domain nor any region in the carboxy-terminal region that would correspond to the IPS-1 transmembrane region were found.

**Figure 1.**
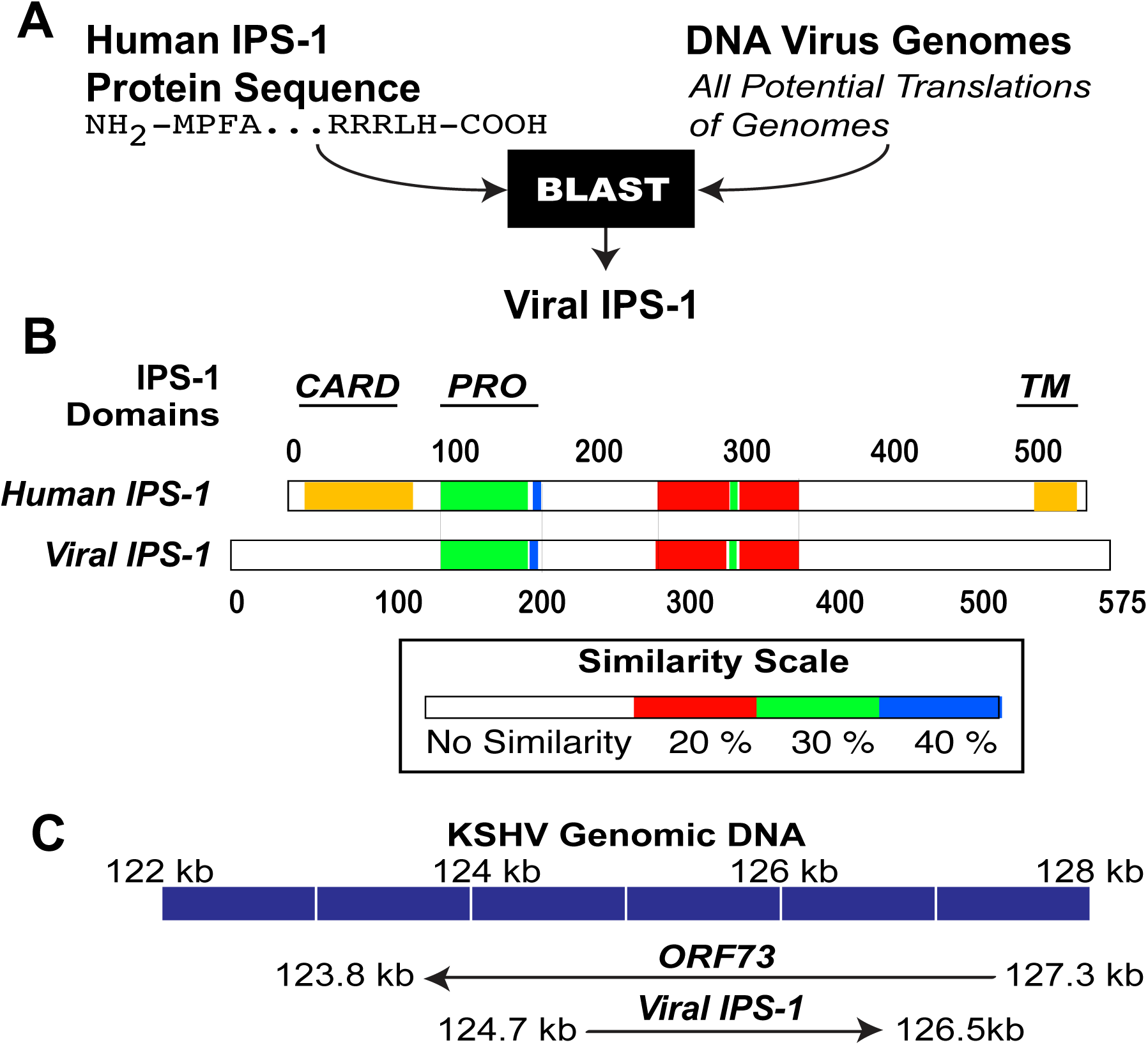
KSHV Encodes a Viral Homologue of IPS-1. (A) To determine possible viral proteins that could target the IPS-1 signaling pathway, we screened the human IPS-1 sequence against all possible translations of the collection of DNA virus genomes. Using BLAST, a search for viral homologues of IPS-1 identified one potential protein after analyzing all possible translations of all DNA virus genomes within the KSHV genome. This novel ORF was termed viral IPS-1 (v-IPS-1). (B) When aligned against the human IPS-1 proteins, v-IPS-1 was found to have homology within the proline-rich (PRO) domain (Green Domain) and an area important for interaction with downstream signaling. A TRAF interacting motif (TIM) is surrounding by a region of low similarity (Red Domain). There was no significant homology with the CARD domain or the transmembrane (TM) domains of human IPS-1 (Both of these are indicated by golden shading in the Human IPS-1 diagram). (C) The novel v-IPS-1 ORF was found to be encoded within the ORF73 (LANA) ORF locus of KSHV, but was in a reverse complement orientation possibly explaining why it was not previously identified.

When mapping this predicted ORF to the genome of KSHV it was found to correspond to a region in the ORF73 locus but is in the reverse complement orientation (Figure 1C).

The v-IPS-1 ORF is smaller than ORF73 with ORF73 and v-IPS-1 being approximately 3.5 kb and 1.8 kb in length respectively. The fact that the v-IPS-1 is in the antisense orientation of ORF73 may explain why this ORF was not yet described.

### The KSHV Viral-IPS1 ORF is Expressed During Lytic Replication

Because the v-IPS-1 ORF was not yet described it was important to develop an antibody to show that we could detect the protein expressed from this ORF during KSHV replication. We had an antibody developed against the C-terminal domain of the v-IPS-1 protein that contained no predicted homology to either KSHV or human proteins (Figure 2A). We also cloned the v-IPS-1 ORF into an expression plasmid and transfected the expression plasmid in HEK 293T cells. Upon immuno-blotting the lysates from the transfected cells we were able to detect a protein of 35 kDa (Figure 2B). This protein is smaller than the predicted molecular weight of the full-length v-IPS-1 ORF protein product. However, based on the position of the antigenic peptide, the region with a predicted TIM would still be intact.

**Figure 2.**
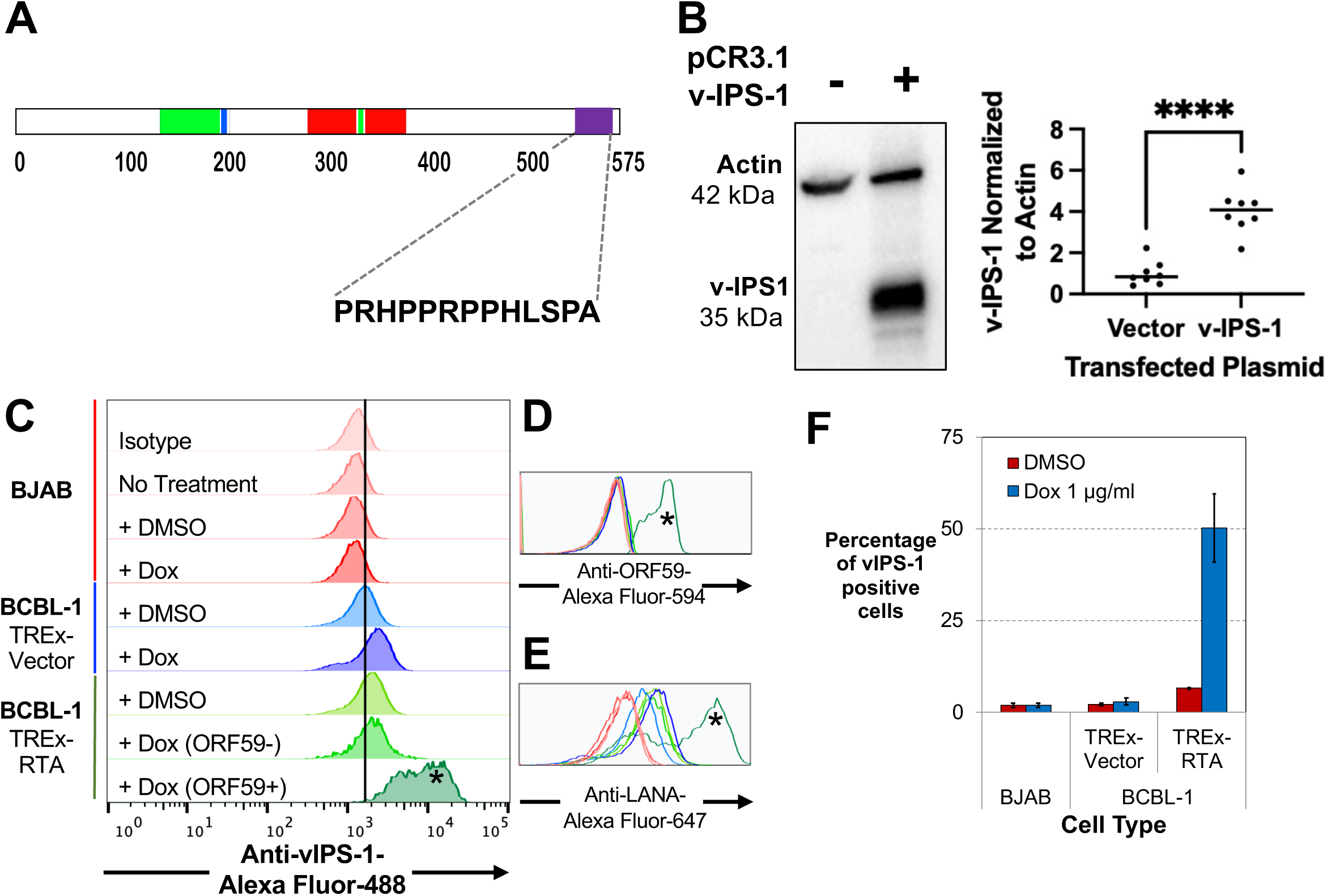
KSHV v-IPS-1 Is a Lytic Gene of KSHV. (A) To determine if v-IPS-1 was indeed a protein produced during KSHV infection, as well as determine where in the latent and lytic parts of the viral replication cycle we developed an antibody was against the unique C terminus of v-IPS-1 as indicated. This amino acid sequences had no similarity to any human or any other KSHV protein. (B) An expression clone of the v-IPS-1 ORF that was PCR amplified from a cDNA library from KSHV infected BCBL1 cells was transfected into HEK 293T cells and whole cells protein lysates were analyzed by western blot using the the anti-v-IPS-1 antibody developed in 2A as well as an anti-Actin antibody. Only cells transfected with the v-IPS-1 expression plasmid showed a band reactive to the v-IPS-1 antibody. To the right of blot is a graph of quantitation of 8 replicates of this experiment with **** standing for p of <0.0001 by unpaired t-test. (C) BCBL-1 cells which hold the latent KSHV genome and were engineered to express RTA under control of a tetracycline inducible promoter (TREx) were analyzed by flow cytometry for the lytic marker ORF59 and the antibody to v-IPS-1 from 2A. BJAB cells, being used as a negative control without KSHV, and BCBL-1 cells with a TREx expressing a vector control as a cell that should not enter lytic phase were also used. Only ORF59+ BCBL-1 cells which are actively in lytic replication express v-IPS-1. (D) Cells in 2C were also stained for ORF59 and the histogram marked with * confirms that high ORF59 expression is in cells expressing v-IPS-1. (E) Cells in 2C were also stained for LANA expression. We confirmed that our BCBL-1 cells do have KSHV latently infected. (F) Three biologic replicates of the flow cytometry shown in 2C were quantitated and the percentage of v-IPS-1 positive cells were graphed. This again confirms that only lytic replicating KSHV infected cells express v-IPS-1.

With an antibody to v-IPS-1 we went on to determine when in KSHV infection the protein is produced. To do that we utilized the BCBL-1 TREx system. These BCBL-1 cells were engineered to express RTA under control of a doxycycline inducible promoter and were used to assess v-IPS-1 expression during KSHV infection [18]. Upon the addition of doxycycline, RTA would be overexpressed inducing the lytic replication program. Using the BJAB cell line as a control we were able to see the background level of staining with the v-IPS-1 antibody (Figure 2C). BCBL-1 TREx, which contain a vector control engineered after the doxycycline inducible promoter and that are still latently infected with KSHV, show no additional antibody staining beyond BJAB staining. However, BCBL-1 TREx cells that are induced to express RTA and begin lytic replication show a 10-fold increase in v-IPS-1 staining in ORF59 positive cells. ORF59 expression can be used as a marker of lytic replication. The v-IPS-1 staining is not seen in ORF59 negative cells strongly implying that v-IPS-1 is expressed when other lytic proteins are expressed. An overlay of the ORF59 and LANA signals for the populations in Figure 2C are shown in Figure 2D and 2E, respectively. The asterisks in Figure 2C, 2D and 2E denote the same cell population. In addition, quantitation over separate experiments shows that consistently approximately 50% of BCBL-1 TREx-RTA cells stimulated with doxycycline express v-IPS-1 (Figure 2F).

### Viral-IPS1 Blocks Interferon-beta Induction at the TRAF3 Signaling Node

The homology of v-IPS-1 to cellular IPS-1 sets up the hypothesis that v-IPS-1 may interfere with the normal function of IPS-1 which is to induce IFN. To test this hypothesis, we used an interferon-beta (IFN-beta) luciferase reporter induced by expression of the various components of the IFN induction pathway (Figure 3A).

**Figure 3.**
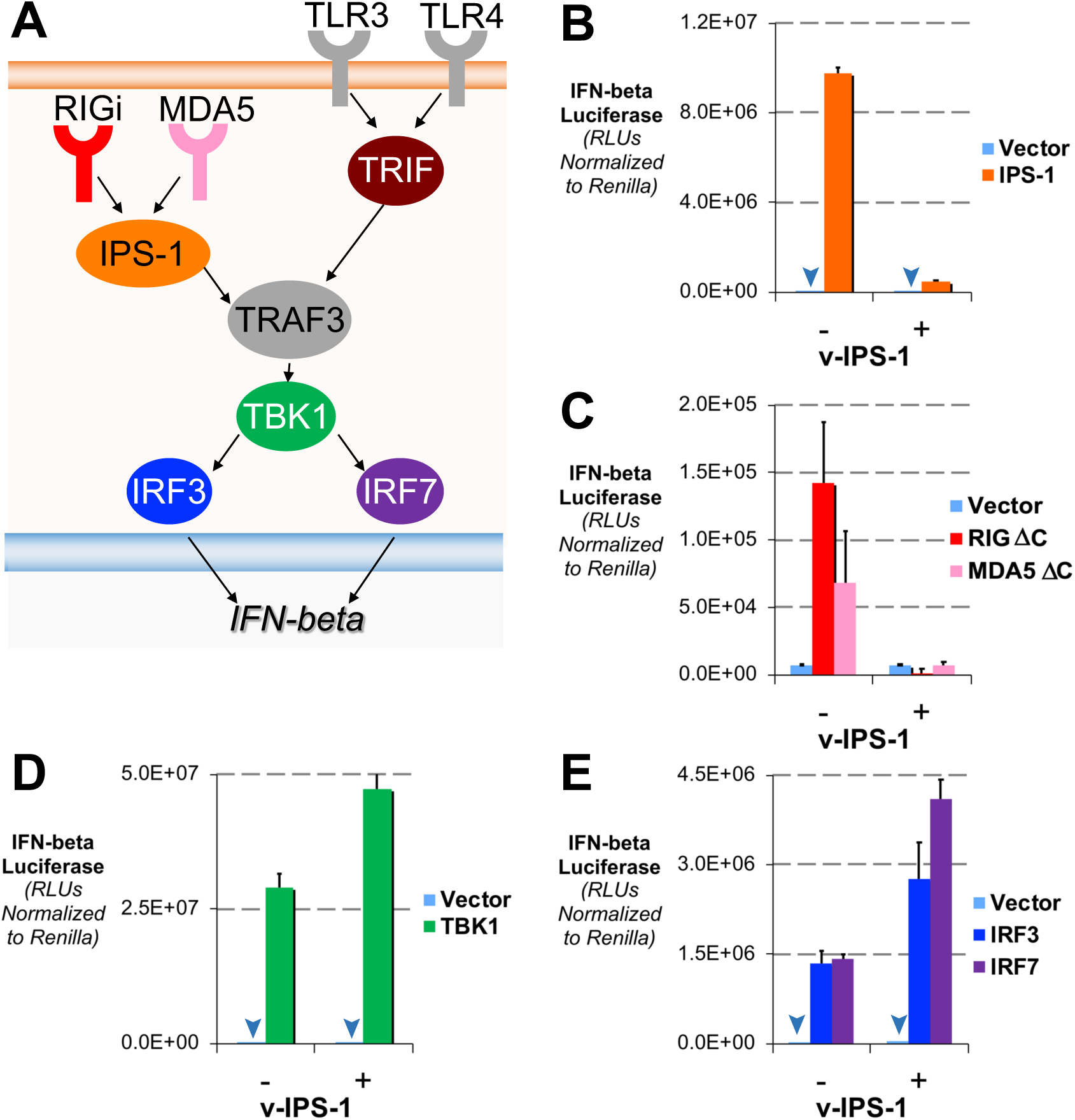
v-IPS-1 Blocks Induction of Interferon Upstream of TRAF3. (A) A diagram representing the interferon induction pathways showing convergence at TRAF3. To determine how v-IPS-1 impacted the different steps of the IFN-beta induction signaling pathway we overexpressed v-IPS-1 along with cellular components of the pathway. (B) HEK 293T cells were transfected with an expression plasmid for IPS-1, which will induce the co-transfected IFN-beta luciferase plasmid which has luciferase under control of the IFN-beta promoter. In addition, these cells were co-transfected with or without the v-IPS-1 expression plasmid as well as a Renilla luciferase reporter. After 18 hours, cells were lysed and firefly and Renilla luciferase was quantitated, with average of triplicates graphed with standard deviations reflected in the error bars. Here v-IPS-1 blocks induction of IFN-beta by IPS-1 overexpression. As in (B), expression plasmids for constitutively active RIG-I and MDA-5 were used in (C), TBK1 in (D) and IRF3 or IRF7 in (E) to see the impact of v-IPS-1 on these different inducers of IFN-beta. Expression of v-IPS-1 blocks interferon induction upstream of TRAF3 when stimulated with either RIG-I, MDA5 or IPS-1. However, v-IPS-1 does not block IFN-beta induction when stimulated by overexpression of TBK1, IRF3 or IRF7, all of which are downstream of TRAF3. Downward arrows are used to mark low values on graphs to point them out in the graphs. All data is representative of technical triplicates and was repeated at least three times.

Expression of cellular IPS-1 induced robust IFN-beta reporter but is blocked by co-expression with v-IPS-1 (Figure 3B). Likewise, expression of either of the RLRs, RIG-I or MDA-5, that are missing their C-terminal domains and thus constitutively active lead to IFN-beta reporter induction that is blocked by expression of v-IPS-1 (Figure 3C).

Induction of IFN-beta reporters by expression of components upstream of TRAF3 are blocked by v-IPS-1. Of note is that exogenous TRAF3 expression induces the IFN-beta reporter minimally and thus there was no statically significant change in reporter activation (Data not shown).

Next, we tested components of the IFN-beta induction pathway that were downstream of TRAF3. Expression of the kinase TBK1 induces strong IFN-beta reporter that is not blocked by co-expression with v-IPS-1 (Figure 3D). Likewise, overexpression of either of the IRFs, IRF3 or IRF7, which induce moderate IFN-beta reporter induction are not blocked by expression of v-IPS-1 (Figure 3E). In summary, v-IPS-1 blocks induction of the IFN-beta reporter by expression of components upstream of TRAF3 but not downstream of TRAF3 (Refer to Figure 3A).

### Expression of Viral-IPS1 Targets the IPS-1:TRAF3 Innate Signaling Node

Because v-IPS-1 expression reduces IFN induction upstream of the TRAF3 signaling node, we next assessed the stability of IPS-1 and TRAF3. Expression of IPS-1 is markedly diminished when co-expressed with v-IPS-1, with 85% decrease in band intensity over multiple experiments (Figure 4A). On the other hand, TRAF3 expression is somewhat stabilized in the presence of v-IPS-1, with 7.3-fold increase in band intensity over multiple experiments (Figure 4B). Because v-IPS-1 is predicted to have an intact TIM, we hypothesized that interaction of v-IPS-1 and TRAF3 leads to both the reduction in IPS-1 expression and increase in TRAF3 stability. To address this possibility, we overexpressed both TRAF3 and IPS-1 in the presence of v-IPS-1 (Figure 4C). We see that only when TRAF3 is also overexpressed do the levels of IPS-1 increase. This strongly implies that interaction of v-IPS-1 with limited cellular TRAF3 leads to changes in IPS-1.

**Figure 4.**
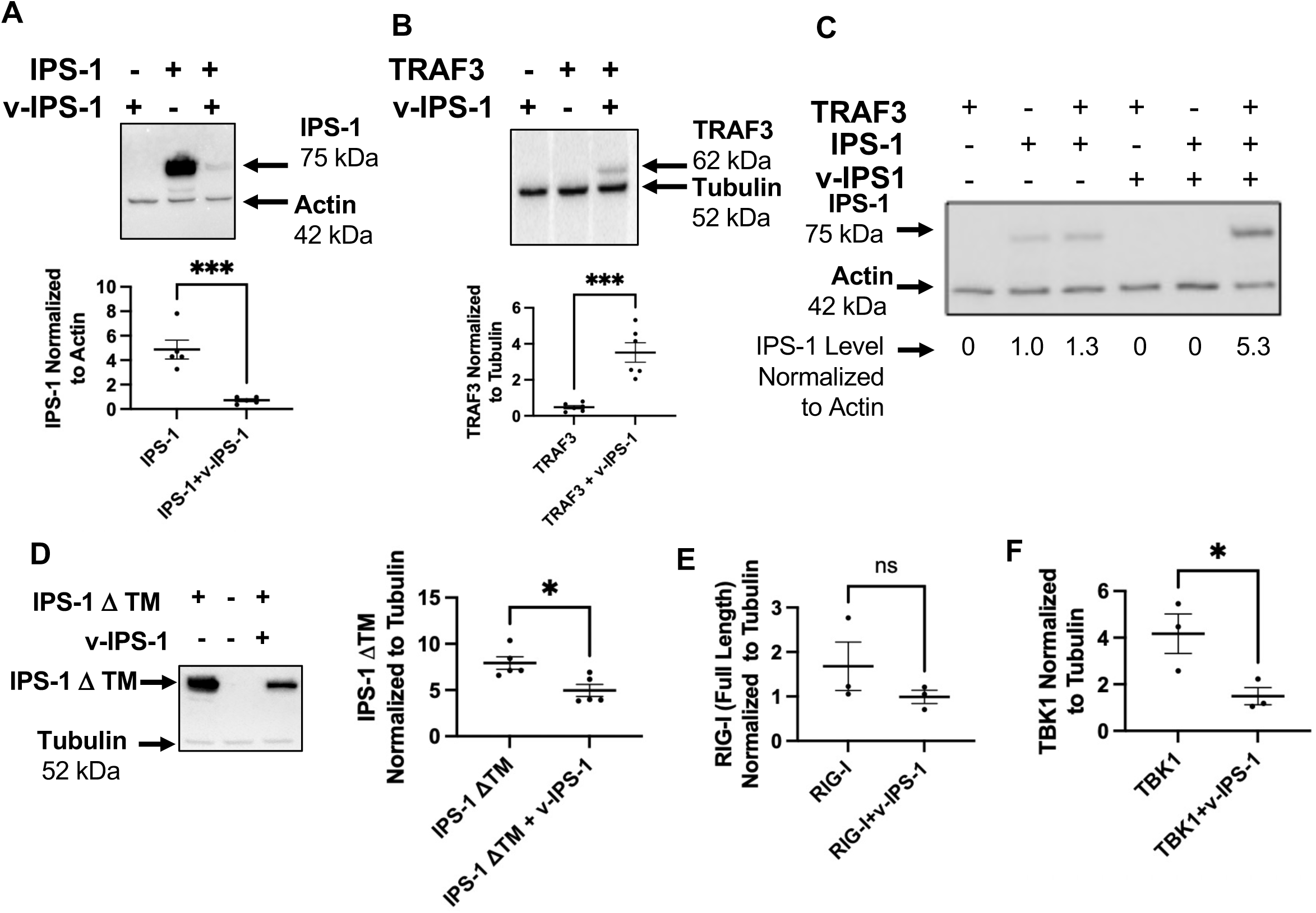
v-IPS-1 Destabilizes IPS-1 while Stabilizing TRAF3. To determine how v-IPS-1 impacts the stability of signaling nodes in the IFN induction pathway, we expressed v-IPS-1 along with different cellular proteins. (A) HEK 293T cells were transfected with an expression plasmid for FLAG-IPS-1 with or without the v-IPS-1 expression plasmid and after 24 hours, cells were lysed for Western blot of FLAG-IPS-1 and Actin. The smaller 52 kDa form of IPS-1 can also be seen in lane 2. Below the blot is a graph of quantitation of 5 replicates of this experiment with *** standing for p of 0.0007 by unpaired t-test. As in (B), expression plasmids HA-TRAF3 were transfected along with or without a v-IPS-1 expression plasmid. After 24 hours, cells were lysed for Western blot of HA-TRAF3 and Tubulin. Below the blot is a graph of quantitation of 5 replicates of this experiment with *** standing for p of 0.0003 by unpaired t-test. (C) Since IPS-1 and TRAF3 often interact, we sought to determine if TRAF3 could overcome v-IPS-1 destabilization of IPS-1. Combinations of HA-TRAF3, FLAG-IPS-1 and v-IPS-1 were transfected into HEK 293T cells and after 24 hours, cells were lysed for Western blotted for FLAG-IPS-1 and Actin. (D) To determine the importance of membrane targeting of IPS-1 on v-IPS-1 destabilization of IPS-1, we used a construct of FLAG-IPS-1 that was missing the transmembrane domain. HEK 293T cells were transfected with FLAG-IPS-1 ΔTM with or without the v-IPS-1 expression plasmid and after 24 hours, cells were lysed for Western blotted for FLAG-IPS-1 ΔTM and Tubulin. To the right of blot is a graph of quantitation of 5 replicates of this experiment with * standing for p of 0.01 by unpaired t-test. (E and F) Similar to the above experiments, Full length FLAG-RIG-I (E) and HA-TBK1 (F) were co-transfected with v-IPS-1 and after 24 hours cells were lysed for Western blotted for FLAG-RIG-I (E) or HA-TBK1 (F) and Tubulin. Graphs of normalized bands whose intensities were quantitated in 3 replicates of this experiment were plotted with ns in (E) standing for no significance and * in (F) standing for p of 0.04 by unpaired t-test.

IPS-1 is localized to membranes, with mitochondrial membranes being the main site of localization of this protein. We used a mutant of IPS-1 that is cytoplasmically localized due to a missing transmembrane domain to determine if IPS-1 localization to the membrane is required for v-IPS-1-mediated degradation. As seen in Figure 4D, expression of IPS-1 that is missing a transmembrane domain is reduced by 36% when co-expressed with v-IPS-1. In comparison, Figure 4A shows that full-length IPS-1 expression is decreased by 85% when co-expressed with v-IPS-1.

### TRIF Signaling is Also Blocked by Viral-IPS1

Besides coordinating RLR signaling, TRAF3 coordinates signaling from other innate immune receptors such as TLR3 and TLR4 (Figure 3A) [19; 20]. To determine if v-IPS-1 also blocks TLR3/4 signaling we expressed TRIF in the presence of v-IPS-1. TRIF expression by western blotting is greatly diminished when co-expressed with v-IPS-1 (Figure 5A). Expression of v-IPS-1 also blocks TRIF-mediated induction of the IFN-beta reporter (Figure 5B). We went on to look at TRAF6 expression which is activated by both IPS-1 and TRIF signaling and saw that TRAF6 expression is lost when co-expressed with v-IPS-1 (Figure 5C) [21; 22].

**Figure 5.**
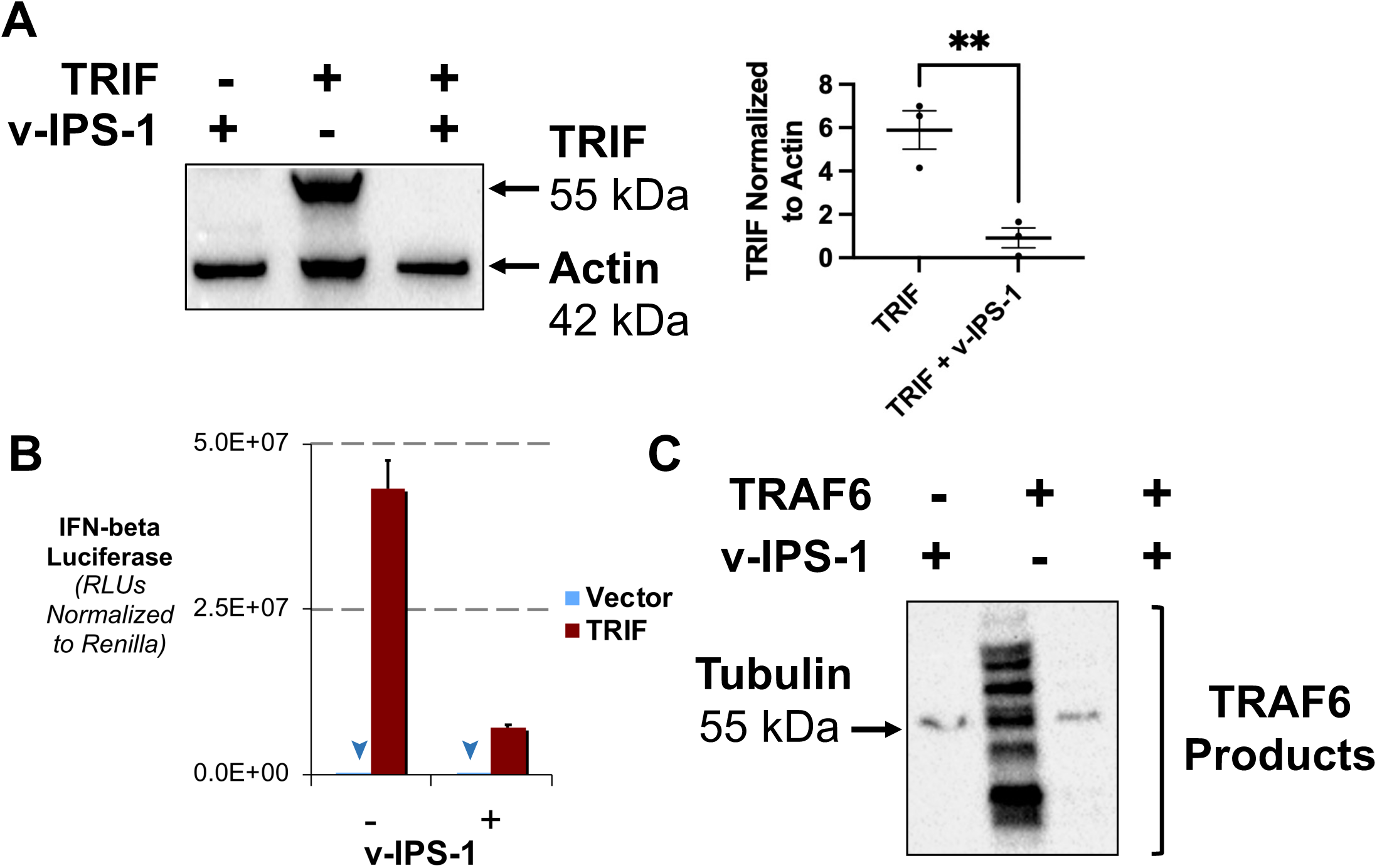
v-IPS-1 Blocks TRIF-mediated Signaling. To determine how v-IPS-1 impacts the TRIF signaling, we expressed v-IPS-1 along with components of TRIF signaling. (A) HEK 293T cells were transfected with an expression plasmid for Myc-tagged with or without the v-IPS-1 expression plasmid and after 24 hours, cells were lysed Western blotted for Myc-TRIF and Actin was done. To the right of blot is a graph of quantitation of 5 replicates of this experiment with ** standing for p of 0.0075 by unpaired t-test. (B) Similar to Figure 3 B-E, HEK 293T cells were transfected with an expression plasmid for Myc-tagged TRIF with or without the v-IPS-1 expression plasmid and after 24 hours, cells were lysed and firefly and Renilla luciferase was quantitated, with average of triplicates graphed. Refer to Figure 3A for diagram of location of TRIF in signaling pathways. Downward arrows are used to mark low values on graphs. (C) HEK 293T cells were transfected with an expression plasmid for HA-tagged TRAF6 with or without the v-IPS-1 expression plasmid and after 24 hours, cells were lysed Western blotted for HA-TRAF6 and Tubulin. The Western blots is representative and part of a set of 3 biological replicates.

### Viral-IPS1 Blocks Interferon Induction but Not Interferon Signaling

Expression of v-IPS-1 leads to blocks to IFN-beta reporter induction as well as destabilization of IPS-1. This led us to assess if v-IPS-1 expression could have consequences on virus replication. To address this, we first transfected cells with v-IPS-1 or IPS-1. As expected, we see that while transfection of IPS-1 strongly blocks replication of VSV that is engineered to express GFP (Figure 6B versus Figure 6A). However, expression of v-IPS-1 leads to strong replication of VSV-GFP (Figure 6C versus 6A). Quantitation of GFP expression by flow cytometry corroborated the microscopy observations showing that IPS-1 stopped almost all VSV replication as monited by GFP but v-IPS-1 allowed for enhanced VSV replication as monitored by GFP (Figure 6D).

**Figure 6.**
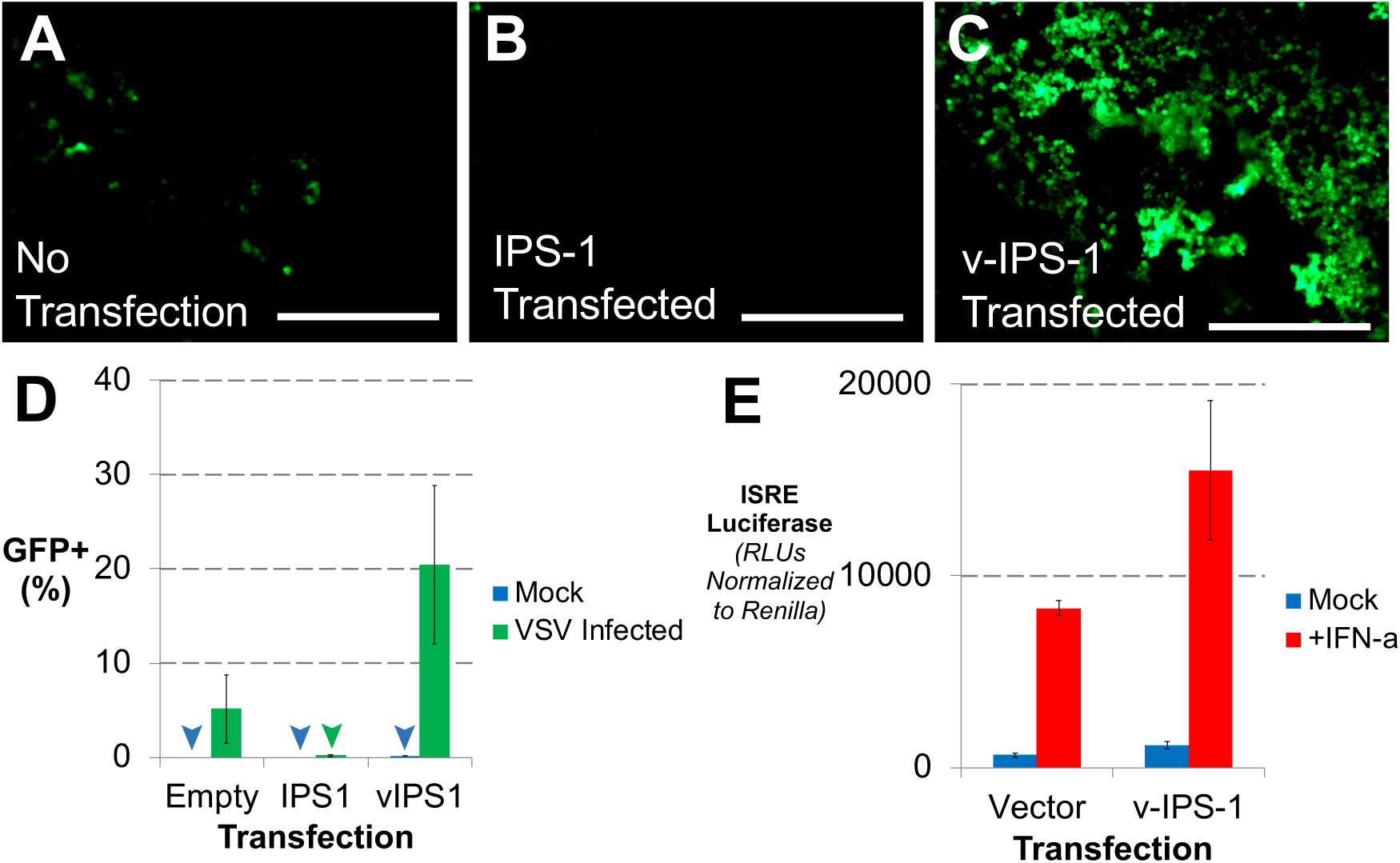
v-IPS-1 Blocks Direct Induction of an Antiviral State. To determine if v-IPS-1 could impact establishment of an antiviral state in cells, we used infection with VSV that was engineered to express GFP. IFN expression blocks VSV replication and in this case should limit GFP expression. HEK 293T cells in A, B and C were infected with VSV-GFP. While cells in (A) were not transfected prior to infection, cells in (B) were transfected with IPS-1 to induce an antiviral state and cells in (C) were transfected with the v-IPS-1 expression plasmids to assess the ability of v-IPS-1 to block antiviral activity during VSV infection. Images of cells were taken, and representative images are shown. The bar represents 400 mm. GFP expression indicated levels of virus replication. (D) Cells from A, B and C were fixed and analyzed by flow cytometry to monitor the percentage of infected cells which is indicated by GFP produced by VSV replication. Downward arrows are used to mark low values on graphs. (E) To determine if v-IPS-1 blocks signaling from external or exogenous IFN, we stimulated v-IPS-1 expressing cells with IFN. HEK 293T cells were transfected with or an empty expression plasmid or one expressing v-IPS-1 expression plasmid cells as well as a firefly luciferase reporter under the control of the Interferon-stimulated response element (ISRE) promoter and Renilla luciferase reporters for 18 hours. ISRE promoters are induced by exogenous IFN stimulation. After an additional 8 hours of stimulation by Interferon-alpha, cells were lysed and firefly and Renilla luciferase was quantitated, with average of triplicates graphed.

The above results strongly imply that there is a disruption in IFN function allowing enhanced virus replication when v-IPS-1 is expressed. While v-IPS-1 does block IFN induction in our earlier data, the impact of v-IPS-1 on signaling induced by signaling from exogenous IFN such as paracrine or pDC released is an importation question.

Expression of v-IPS-1 does not block signaling after cells are stimulated with exogenous interferon (Figure 6E). This focuses v-IPS-1 function on its impact on IFN induction which is critical in the antiviral responses induced by KSHV infected cells.

## DISCUSSION

Induction of innate immunity is mediated by diverse families of Pattern Recognition Receptors (PRRs) that recognize molecular “signatures” of invading microbial pathogens termed pathogen associated molecular patterns (PAMPs). Most nucleated cells are capable of sensing and responding to replicating viruses within their cytoplasm via intracellular receptors of the RIG-I-like receptor (RLR) family, which recognize replicating viral RNA structures [23; 24; 25; 26; 27; 28; 29; 30]. RLR receptors signal via cellular IPS-1 to induce IFN production towards development of an antiviral state to limit virus growth. However, viral pathogens need to limit induction of IFN to better replicate.

We used bioinformatics to identify a new viral ORF within the KSHV genome we have termed viral IPS-1 due to its homology to the cellular IPS-1 protein, a key adaptor in detection of cytoplasmic viral nucleic acids. KSHV has several viral encoded genes that are homologues of cellular proteins including viral versions of IL-6, cyclin D, IRFs, and a GPCR[31; 32; 33; 34]. These types of proteins modulate cellular pathways related to the cellular function of the host protein in an uninfected state. Most of these proteins were found by direct genome annotation, however, the lack of identification of v-IPS-1 may be due to its reverse complement orientation within the ORF73/LANA ORF as many KSHV genome annotations exist and while none have identified a transcript for v-IPS-1, there are potential transcription start sites that may cover this region[35].

Expression of the v-IPS-1 has functional consequence of disrupting induction of IFN with stimulation from signaling above TRAF3. Cellular IPS-1 interacts with TRAF3 to coordinate activation of kinases that can then activate the cytoplasmically localized IRF transcription factors that translocate to the nucleus upon phosphorylation to induce transcription of IFN. Signaling by the RLRs (RIG-I or MDA5), cellular IPS-1 or TRIF (a TLR3/4 adaptor also upstream of TRAF3 [36]) is blocked by v-IPS-1 (Figure 3 and 5), while neither the kinase TBK1 nor the transcription factors IRF3 and IRF7 are affected by v-IPS-1 (Figure 3D and 2E). This data shows that v-IPS-1 modulates at signaling at or upstream of TRAF3, a major signaling adaptor protein that links the upstream receptor/adaptor signaling with the downstream kinase/transcription factor steps in IFN induction.

With these results in mind, it is important to note that the strongest evidence of interaction between TRAF3 and cellular IPS-1 was after stimulation by viral PAMPs inducing a detectable interaction [17]. Without IPS-1, IFN levels are not sufficient to control herpesvirus infections [37]. Conversely, TRAF3 deficiency leads to enhanced IFN release in herpesvirus infections [38]. In addition, there is upregulation in induction of IFN pathway by blocking the TRAF3 ubiquitination pathways strongly implying that the presence of TRAF3 is a negative modulator of IFN signaling [39]. Overall, v-IPS-1 disruption of the TRAF3 and IPS-1 interactions fits with the biology of theTRAF3/IPS-1 axis and v-IPS-1 could be taking advantage of this axis to disrupt IFN induction. Future determination of possible direct or indirect interactions between v-IPS-1 and TRAF3 or IPS-1 is important to fully explore the mechanism of this protein.

However, the exact impact of v-IPS-1 on KSHV pathogenesis is not a straightforward question as KSHV encodes several other IFN antagonizing proteins including the viral-IRF proteins and LANA [6; 7; 8; 9; 10]. While Figure 6E shows that v-IPS-1 does not block signaling by exogenous IFN, ORF10 is known to limit IFN signaling toward an antiviral state through the IFNAR receptor [40]. Another question is how important is the IPS-1 pathway in signaling for a herpesvirus as canonically the primary herpesviral PAMPs are thought to focus on the herpesviral DNA and not viral RNA products.

Detection of cellular RNAs by the RLR pathway may be important in inducing IFN by KSHV infection [41]. This process is important in limiting virus replication as depleting IPS-1 prior to lytic reactivation increases the levels of virus output [42]. It may be that beyond purely blocking IFN, v-IPS-1 may block TRAF3 signaling or other TRAF proteins thus disrupting signaling from other receptors. For example, the role of TRAF3 in signaling in B cells and even lymphoma development highlights the importance of determining the full set of outcomes induced by TRAF3 modulation by v-IPS-1 [43]. Future work into the role of v-IPS-1 during active infection may also show a dynamic of the v-IPS-1 and overlapping LANA/ORF73 locus.

This study has shown important of the v-IPS-1 protein in modulating innate immune signaling. An important limitation of our study is that to determine the molecular basis of this protein, we have had to study it in isolation from other viral proteins. The relative importance of v-IPS-1 in comparison to other modulators of innate immune signaling such as vIRFs cannot be determined our study. Future studies into the relative contribution of v-IPS-1 in comparison to other modulators of innate immune signaling are ongoing. An additional limitation is that this paper does not address how v-IPS-1 impacts IFN induction during viral infection is difficult to discern from our studies for the same reason. We are actively working on this topic. Finally, we have not yet determined how v-IPS-1 interacts with either IPS-1 or TRAF3. Determining if there is direct interaction or indirect connections between v-IPS-1 and these proteins is critical to fully elucidating the mechanism of this viral protein.

## CONCLUSION

Here we report the discovery of a KSHV homologue of IPS-1, a key protein in the induction of IFN in response to viral infection. This v-IPS-1 protein blocks IFN induction from both the RLR and TLR pathways and leads to destabilization of IPS-1 and concatenate stabilization of TRAF3. The v-IPS-1 protein also seems to specifically block the establishment of the antiviral state during infection. This is the first viral homologue of IPS-1 and underscores the importance of IPS-1 in establishing viral infection.

## METHODS

### Cell Lines Used

Human embryonic kidney 293T cells (ATCC Catalogue number were grown in Dulbeco’s Modified Eagle Medium (DMEM, high glucose) from Thermo Fisher. Media for HEK 293T is supplemented with 5% fetal bovine serum (FBS) and 1% penicillin/streptomycin (P/S) antibiotics. Cells were plated in TC treated dishes (150X20mm from Genesee Scientific) and kept in the incubator at 37℃ and 5% CO_2_. Cells were passaged every 2 days. BJAB cells were grown in Roswell Park Memorial Institute (RPMI) 1640 Medium from Thermo Fisher supplemented with 10% fetal bovine serum (FBS) and 1% penicillin/streptomycin (P/S) antibiotics. BCBL cells (TREx-Vector and TREx-RTA) were grown as previously described and used as originally described by the Jung group [18].

### Luciferase reporter assay

Luciferase reporter assay was performed to measure the Interferon-β promoter activity. HEK 293T cells were transfected in 24-well dishes as sets of triplicates. In brief, triplicates were transfected with X-tremeGene at a ratio of 2:3 (transfection reagent: DNA). Each triplicate was transfected 1250 ng of total plasmids with a mix of 200 ng of pGL3-Interferon-β luciferase reporter, 50 ng of pSV40-Renilla luciferase reporter and 1000 ng of expression plasmid. Figure 6E used pGL3 which has ISRE instead of the Interferon-β promoter. Transfection complex was incubated in 75 μl of DMEM with X-tremeGene per triplicate for 20 mins. HEK 293T cells were plated at 1x10^5^ cells total per well. To each well 20 μl of transfection complex were added. Transfections were incubated for 24 hours. The next day, cells were lysed with passive lysis buffer from the Dual-Luciferase Reporter Assay kit (Promega). Luciferase readings were obtained by the GloMax®-Multi Detection System.

### Protein Co-expression Analysis

DNA expression plasmids of human IPS-1, as well as other components of RLR signaling pathway including TRAF3, RIG-I, TBK1, and TRIF, were co-transfected with v-IPS-1 in HEK 293T cells grown in 6-well plates. In our experiments, human FLAG-tagged IPS-1 (1μg), HA-tagged TRAF3 (1μg), FLAG-tagged RIG-I (1μg) or Myc-Tagged TRIF (1μg) were co-transfected with equal amount of the v-IPS-1 (1μg) plasmid. Human FLAG-tagged IPS-1 (1μg), HA-tagged TRAF3 (1μg), FLAG-tagged RIG-I (1μg) and Myc-tagged TRIF (1μg) were also co-transfected with equal amount of GFP plasmid (1μg) as the control group. Cells were transfected at a density of 2.5X10^6^ cells per 6 well dish using Xtreme Gene HP from Sigma-Aldrich.

After transfecting cells for 48 hours, media is aspirated off and cells were lysed by adding 200μl 1X SDS lysis buffer directly onto cells. Cells were scraped from the bottom of the wells and passed through a 21G (0.8mm X 25mm) needle in order to break apart the cell membranes. Samples were loaded into a 4–20% Mini-PROTEAN® precast polyacrylamide gel (Bio-Rad), and gel electrophoresis was run in a Mini-PROTEAN® Tetra cell (Bio-Rad) for 30 minutes at 200V in 1X SDS running buffer (Bio-Rad). After proteins were separated, the proteins are transferred from the gel to a PVDF membrane using the Trans-Blot Turbo Transfer System (Bio-Rad), using the pre-programmed turbo transfer program. Membranes were blocked for 1 hour in 25mL of a blocking solution made by 5% powdered, non-fat milk in 1X TBS with 0.1% Tween 20. Solution containing primary antibodies for FLAG-tagged proteins are made by adding Anti-FLAG M2 antibodies (produced in mouse) from SIGMA into 10mL of the blocking solution (1:10000). Solution containing primary antibodies for Myc-tagged proteins are made by adding Myc-Tag 71D10 antibodies (produced in rabbit) from Cell signaling into 10mL of the blocking solution (1:10000). Solution containing primary antibodies for HA-tagged proteins are made by adding Anti-HA HA-7 antibodies (produced in mouse) from SIGMA into 10mL of the blocking solution (1:20000). Primary antibody solution against v-IPS-1 is made by adding 10μL of a rabbit produced antibody against v-IPS-1’s C-terminal sequence (1:1000) that was produced by NeoBioLab. Tubulin is blotted by primary antibody solution of beta-Tubulin 9F3 antibodies (produced in rabbit) from Cell signaling in 10mL of the blocking solution (1:10000). Actin is also blotted by primary antibody solution of beta-Actin 8H10D10 antibodies (produced in mouse) from Cell signaling in blocking solution (1:10000). After 1 hour the blocking solution is replaced by blocking solution with primary antibodies and incubated at 4℃ refrigeration overnight.

After 24 hours the solution with primary antibodies was removed washed, and secondary antibodies (goat anti-mouse lgG-HRP sc-2005 from Santa Cruz against membranes blocked by Anti-FLAG, Anti-HA and beta-Actin, goat anti-rabbit lgG-HRP sc-2004 from Santa Cruz against membranes blocked by Myc-tagged antibody, Tubulin and v-IPS-1) and incubated on rocker in room temperature for 1 hour. Afterwards, solution with secondary antibodies was washed extensively. A mixed luminescent solution made from the ImmunoCruz Western Blotting Luminol Reagent (SantaCruz sc-2048) was added to the membrane and incubated for 1 minute at room temperature before imaging in the Image Lab software on a ChemiDoc MP Imaging System machine (Bio-Rad) . Western blot ladders are imaged in Blot Ladder no bin program and target proteins are imaged in Blots Chemi program.

Band intensity was determined with the Image Lab software on a ChemiDoc MP Imaging System. Bands were normalized using Microsoft Excel with target bands divided by housekeeping bands (Actin or Tubulin). Normalized bands were further analyzed in GraphPad Prism 10 and statistical differences in groups of experiments were analyzed by unpaired t-test and graphed.

### Flow Cytometry Analysis for Protein Expression

Antibodies against ORF59 (ABI), LANA and v-IPS-1 were used to stain BJAB and BCBL-1 cells. Cell protocol was adapted from protocol provided by Cell Signaling Technology. In brief, cells were collected by centrifugation and media was removed and washed with 1x PBS. Cells were resuspended in approximately 100 µl of 4% formaldehyde and allowed to incubate and fix for 15 min at room temperature. Cells were permeabilized by adding ice-cold 100% methanol slowly to cells, while gently vortexing, to a final concentration of 90% methanol. Cells were allowed to incubate and permeabilize for 10 minutes on ice. Cells were washed with 1x PBS to remove methanol.

100 µl of primary antibody diluted in 1x PBS + 0.5% BSA was added to cell pellets and incubated for 1 hour at room temperature. Pellet was washed with 1x PBS + 0.5% BSA before resuspending cells in 100 µl of diluted fluorochrome-conjugated secondary antibody. Cells were incubated for 30 minutes at room temperature. Cell pellets were washed with 1x PBS + 0.5% BSA and cells were resuspended cells in 500 µl of 1X PBS and analyzed using an BD Accuri™ C6 Flow Cytometer (BD Biosciences).

### Interferon Bioassay

HEK 293T cells prepared as described above, were grown to 80% confluency. Cells were trypsinized, counted and diluted to the concentration of 2.5X10^6^ cells for a 6-well plate. IPS-1 and v-IPS-1 plasmids were transfected into cells using Xtreme Gene HP from Sigma-Aldrich. After transfection, cells were incubated for 30 hours at 37℃. VSV-GFP is an interferon-sensitive virus that was engineered to encode GFP [44]. VSV-GFP infection solution was made by adding 1 μl of VSV-GFP (kept at 5x10^4^ PFU/μL) into 600μL non-supplemented DMEM media and mixed to homogeneity by vortexing for 5 seconds. For each treatment group, 48μL of the VSV-GFP infection solution was added to 3 wells of the total 6 wells. Cells were further incubated for 18 hours at 37℃.

Cells from each well were resuspended with a pipette into a microcentrifuge tube and pelleted at 3000 rpm for 5 minutes. After spinning down, DMEM media was pipetted out and cells were fixed by adding 1% PFA in 1X PBS and vortexing to mix. GFP positive cells were measured by flow-cytometry using an BD Accuri™ C6 Flow Cytometer (BD Biosciences).

## ACKNOWLEDGEMENTS

Research was supported by NIH Grants R15AI138847, R03DE023307 and R21CA261297 to DJS. The granting agency had no role in overall study design, the design, analysis, and interpretation of the data, decision to publish, or preparation of the manuscript.

## AUTHOR CONTRIBUTIONS

DJS and DMJr were involved in the conception of the project. DMJr, BH, JCS, AS, JRP, JTT and DJS were involved in the design, analysis, and interpretation of the data. DJS, DMJr, and BH were involved in drafting of the paper with contributions from all authors for experimental detail, while DJS and DMJr were responsible for revising it critically for intellectual content. All authors provided final approval of the version to be published; In addition, all authors agree to be accountable for all aspects of the work.

## DATA AVAILABILITY STATEMENT

The data that support the findings of this study are available from the corresponding author, DJS, upon reasonable request. Additional data for this paper is freely available at DOI 10.17605/OSF.IO/PFR94 or at https://osf.io/pfr94/?view_only=3164b9a33c2c4bd6bba8864011e8e1be.

## DECLARATION OF INTERESTS

The authors declare no competing interests.

## Notes

### Competing Interest Statement

The authors have declared no competing interest.

